# Integrative machine learning approaches for predicting disease risk using multi-omics data from the UK Biobank

**DOI:** 10.1101/2024.04.16.589819

**Authors:** Oscar Aguilar, Cheng Chang, Elsa Bismuth, Manuel A. Rivas

**Author notes:** Contributing authors.

## Abstract

We train prediction and survival models using multi-omics data for disease risk identification and stratification. Existing work on disease prediction focuses on risk analysis using datasets of individual data types (metabolomic, genomics, demographic), while our study creates an integrated model for disease risk assessment. We compare machine learning models such as Lasso Regression, Multi-Layer Perceptron, XG Boost, and ADA Boost to analyze multi-omics data, incorporating ROC-AUC score comparisons for various diseases and feature combinations. Additionally, we train Cox proportional hazard models for each disease to perform survival analysis. Although the integration of multi-omics data significantly improves risk prediction for 8 diseases, we find that the contribution of metabolomic data is marginal when compared to standard demographic, genetic, and biomarker features. Nonetheless, we see that metabolomics is a useful replacement for the standard biomarker panel when it is not readily available.

## 1 Introduction

Chronic diseases constitute the primary causes of mortality and morbidity in the United States. Numerous research efforts have been dedicated to the prediction of chronic disease risks, with a focus on genomics, lifestyle, nutrition, and demographics, aiming to detect, prevent, and treat these conditions at an earlier stage. It is unclear how these data types perform for disease risk assessment.

In this study, we focus on training disease risk prediction models using four types of data: demographics, genomics, metabolomics, and clinical biomarkers. While demographic (e.g. age and sex) and biomarker (e.g. glucose or cholesterol level in blood samples) data are commonly used to diagnose diseases in clinical settings, other types of data with greater complexity may reveal more information about potential disease onset for patients [1]. Genetic data contains information about the heritable component of a disease from birth. Therefore, they may contain important information about potential disease onsets [2]. Additionally, metabolomic data, the large-scale study of metabolites in the human body, may offer additive values of clinical and demographic predictors, enhancing the predictive power across multiple diseases [3].

Traditional models for disease prediction have primarily focused on singular data types, often overlooking the complex interplay between various biological and demographic factors. Recent studies have shown the potential of individually using metabolomics, demographics, and genomics to predict age-related diseases and mortality [4]. These findings underscore the potential of using machine learning models to leverage high-throughput data and better understand the complex non-linear relationships between multi-omics predictors. In addition, machine learning methods offer a new way of integrating multi-omics data using different fusion methods that we will explore further [5].

## 2 Results

A Multi-omic Risk Score (MRS) was developed for 22 diseases with at least 1000 cases in the UK Biobank, using a variety of binary classification models. In addition, survival models were trained for a subset of diseases using Cox proportional hazard models with L1 penalty, using age-of-onset data from the UK Biobank. Finally, results were integrated into the Global Biobank Engine [6] for public access.

### Binary Classification Models

To predict the onset and diagnosis of disease within the 13-year period following data collection, we trained models on the combined demographic, genomics, metabolomic, and biomarker feature space. We focused on four classifiers: ADA Boost and XG Boost (tree-based ensemble models), Lasso regression (logistic regression with L1 penalty), and MultiLayer-Perceptron (a relatively small Artificial Neural Network (ANN)). After tuning the appropriate hyperparameters for each model, we found that model performance for each classifier was fairly similar across the many diseases. Table 1 provides a full breakdown of model performance across our four major classifiers: ADA Boost, XG Boost, Lasso regression, and Multi-Layer Perceptron. In general, we found XG Boost to be the quickest to train; ADA Boost to provide the sparsest feature selection; and Lasso regression to provide the best classification performance. The average performance of Lasso was an AUC of 0.739 versus 0.727 for XG Boost, 0.726 for ADA Boost, and 0.675 for MultiLayer-Perceptron.

**Table 1:** Comparison of the test AUC for various binary classification models trained on the full feature set. The first four columns list the performance (from left to right) of the Multi-Layer Perceptron, XG Boost, ADA Boost, and Lasso Regression classifiers, with bolded values denoting the highest AUC for each disease. The last three columns show the performance from using logistic regression with different regularization penalties: Ridge (*l*_2_), ElasticNet [7], and None (unregularized). Bolded values indicate improvements over Lasso (*l*_1_ regularization).

| Model | MLP | ADAB | XGB | Lasso | Ridge | Elstic | None |
| --- | --- | --- | --- | --- | --- | --- | --- |
| arthritis (nos) | 0.654 | 0.650 | 0.640 | **0.663** | **0.667** | 0.661 | 0.663 |
| asthma | 0.628 | **0.632** | 0.630 | **0.632** | **0.638** | **0.637** | **0.635** |
| atrial flutter | **0.783** | 0.768 | 0.764 | 0.779 | 0.779 | 0.777 | 0.777 |
| cellulitis | 0.501 | 0.608 | 0.610 | **0.611** | 0.611 | 0.604 | **0.612** |
| cholelithiasis | 0.722 | 0.707 | 0.712 | **0.726** | **0.727** | 0.726 | 0.726 |
| chronic renal failure | 0.867 | 0.864 | 0.866 | **0.873** | **0.875** | 0.871 | **0.874** |
| cystitis | 0.659 | 0.673 | **0.680** | 0.678 | **0.684** | **0.683** | **0.680** |
| diabetes | 0.500 | 0.915 | **0.921** | **0.921** | **0.922** | 0.921 | 0.921 |
| female genital cancer | 0.751 | 0.772 | 0.780 | **0.781** | 0.775 | 0.778 | 0.781 |
| gall stones | 0.671 | 0.713 | 0.708 | **0.724** | **0.725** | **0.725** | **0.725** |
| glaucoma | 0.549 | 0.680 | 0.676 | **0.687** | 0.682 | 0.683 | 0.686 |
| gout | 0.865 | 0.860 | 0.873 | **0.875** | 0.875 | 0.871 | 0.874 |
| irritable bowel syn. | 0.596 | 0.609 | 0.608 | **0.624** | **0.626** | 0.621 | 0.624 |
| male genital cancer | 0.796 | 0.842 | **0.853** | 0.852 | 0.852 | 0.852 | 0.852 |
| myocardial infarction | **0.825** | 0.802 | 0.800 | 0.816 | 0.816 | 0.816 | 0.816 |
| non-mel. skin cancer | 0.500 | 0.694 | 0.701 | **0.715** | **0.716** | **0.716** | **0.716** |
| osteoarthritis | 0.614 | 0.695 | 0.680 | **0.698** | 0.698 | 0.697 | 0.698 |
| periph. vasc. disease | 0.702 | 0.697 | 0.710 | **0.725** | 0.725 | 0.720 | 0.725 |
| psoriasis | 0.503 | 0.647 | 0.658 | **0.679** | **0.683** | 0.679 | **0.680** |
| renal failure | 0.817 | 0.813 | 0.816 | **0.829** | 0.829 | 0.828 | 0.829 |
| skin cancer | 0.702 | 0.686 | 0.691 | **0.704** | **0.705** | 0.704 | **0.705** |
| ulcerative colitis | 0.656 | 0.638 | 0.613 | **0.664** | **0.672** | **0.674** | **0.670** |
| Average Test AUC | 0.675 | 0.726 | 0.727 | **0.739** | **0.740** | 0.738 | **0.740** |

Next, we explored the use of different penalties for our logistic regression models, with results also in Table 1. In general, *LogisticRegressionCV* produced models with between 50 and 150 non-zero coefficients whereas most of our ADA Boost models had around 5 - 30 non-zero features (Figure A1). While both models tend to agree with on the 5-10 most important features, certain differences emerge when comparing relative feature rankings between the two models (Figure 1). When predicting diabetes for instance, both models agreed on five of the top six predictors but differed on the importance of glucose versus creatinine. For myocardial infarction, ADA Boost and Lasso select most highly entirely different sets of metabolites. Rather than linoleic acid or triglycerides, ADA Boost selects most highly for Cholesteryl esters in IDL (top right of Figure 1).

**Fig. 1:**
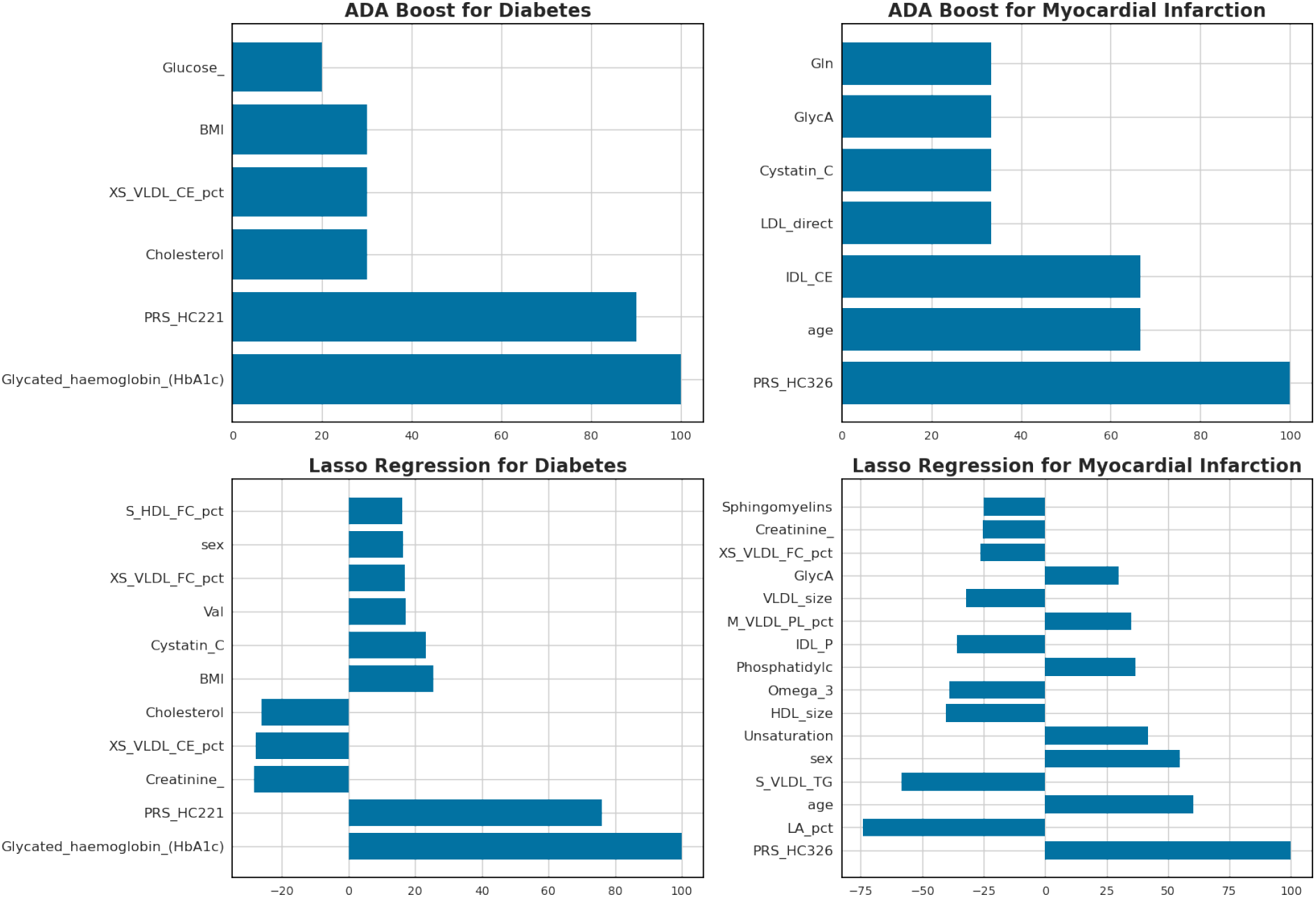
Feature importance from ADA Boost and Lasso for prediction of diabetes and myocardial infarction. The plots are limited to the subset of features from the diabetes and MI models with relative importance ≥ 15 and 25, respectively. Relative importance is displayed as percentage of the strongest feature component.

To assess the contributions of each distinct dataset to the effectiveness of our models, we trained our best-performing models on different combinations of the feature types. By comparing their test set AUC in Figure 2, we observe interesting patterns. The Lasso models for myocardial infarction, for example, did not increase in performance when adding genomics data to the base set of demographic features (AUC of 0.76 for both). The largest difference in AUC (delta of 0.05) came from incorporating common biomarkers in our model, with Cholesterol and Apolipoprotein B selected almost as highly as age and sex. Finally, AUC marginally increased to 0.83 (delta of 0.02) upon adding the full set of metabolomic features, with the ratio of linoleic acid to total fatty acids and the level of triglycerides in small VLDL ranked the highest. Both improvements in AUC were statistically significant, however, with p-values less than 0.0001 when comparing ROC curves with the prior model (Table A2).

**Fig. 2:**
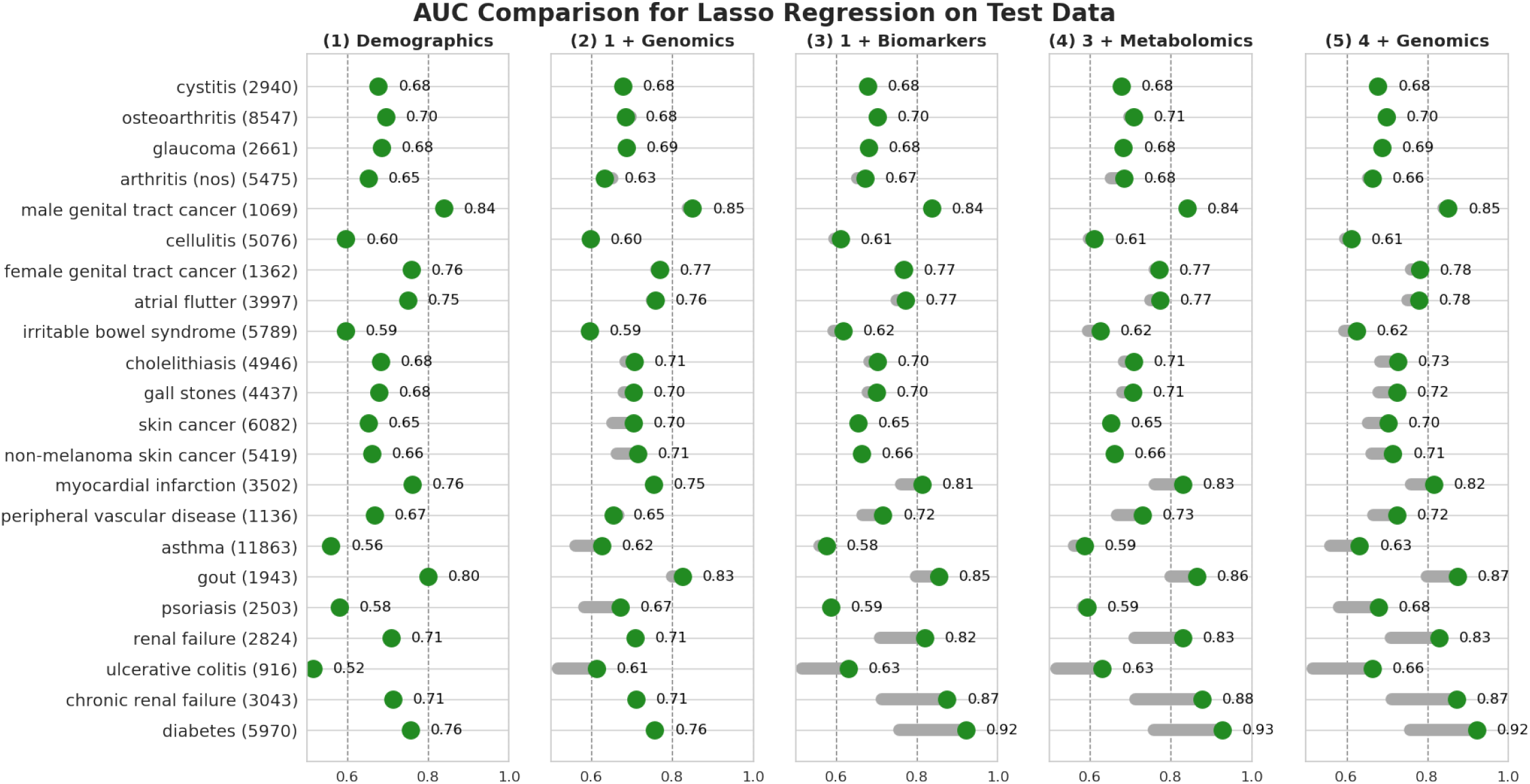
Comparison of the test AUC of lasso regression models trained on different subsets of the data. Green dots represent the AUC (area under the ROC curve) on the test set for each disease, achieved by a model trained on the indicated group of features. Solid bars represent the change in performance from the baseline demographics model (first column). The numbers in the parentheses by the disease names are the number of cases for each disease in the dataset.

The contributions across all diseases of the Genomics and Biomarker data to the predictive performance of baseline demographic models are shown in columns 2 and 3 of Figure 2, respectively. For a few diseases a strong genetic component can be identified. Model test set AUC increased by 0.05 for Non-melanoma skin cancer, 0.06 for asthma, and 0.08 for psoriasis with the addition of genomics data. For many other diseases, adding just the 35 biomarkers yields a significant improvement in performance. The test AUC of the models for peripheral vascular disease, renal failure, chronic renal failure, and diabetes each increased by between 0.05 and 0.16 upon the incorporation of biomarker data. The p-values for the improvement in these ROC curves over the demographic models are all less than 0.0005.

Column 4 of Figure 2 shows the results of adding the full set of 249 metabolomic features to the biomarkers and demographic features. Of the fifteen models which significantly improved with the biomarker data (p *<* 0.05), nine further improve their performance with p *<* 0.05 upon adding metabolomic data (Table A2). The corresponding delta AUC between column 3 and 4 is 0.01 - 0.02 for diabetes, renal failure, chronic renal failure, peripheral vascular disease, myocardial infarction, and arthiritis. The final column in Figure 2 shows the results of incorporating a full feature space into the Lasso model, with the significance tests in column 4 of Table A2 evaluating the minimum improvement in AUC over all previous models. Only the final models for asthma, gout, gall stones, cholelithiasis, and ulcerative colititis had a statistically significant improvement in performance (p *<* 0.05) when comparing ROC curves with prior models. The delta AUC for these models ranged from 0.005 to 0.03, while all others saw virtually no improvement (delta AUC *<* 0.005) over prior models. Furthermore, the diseases with the largest increases in performance from the multi-omics model have both genomic and metabolomic components. Lasso models for ulcerative colitis, most notably, gain 0.09 and 0.11 AUC when adding genomics and biomarker/metabolomic data separately versus 0.14 AUC when used jointly.

### Survival Analysis Models

We train a Cox proportional hazard model for each disease of interest to evaluate the changing risk of patients through time. We then use the Cox models to generate the risk trajectories of individuals within the dataset. Similar to the feature type analysis for the binary classification models, we compare the contributions of the demographic, biomarker, genomics, and metabolomics features with the concordance index (C-index) of the models. Using models with only demographic data as the baseline, we incrementally train models with additional genomics, biomarker, and metabolomics features. We now present the results as listed in Figure 3.

**Fig. 3:**
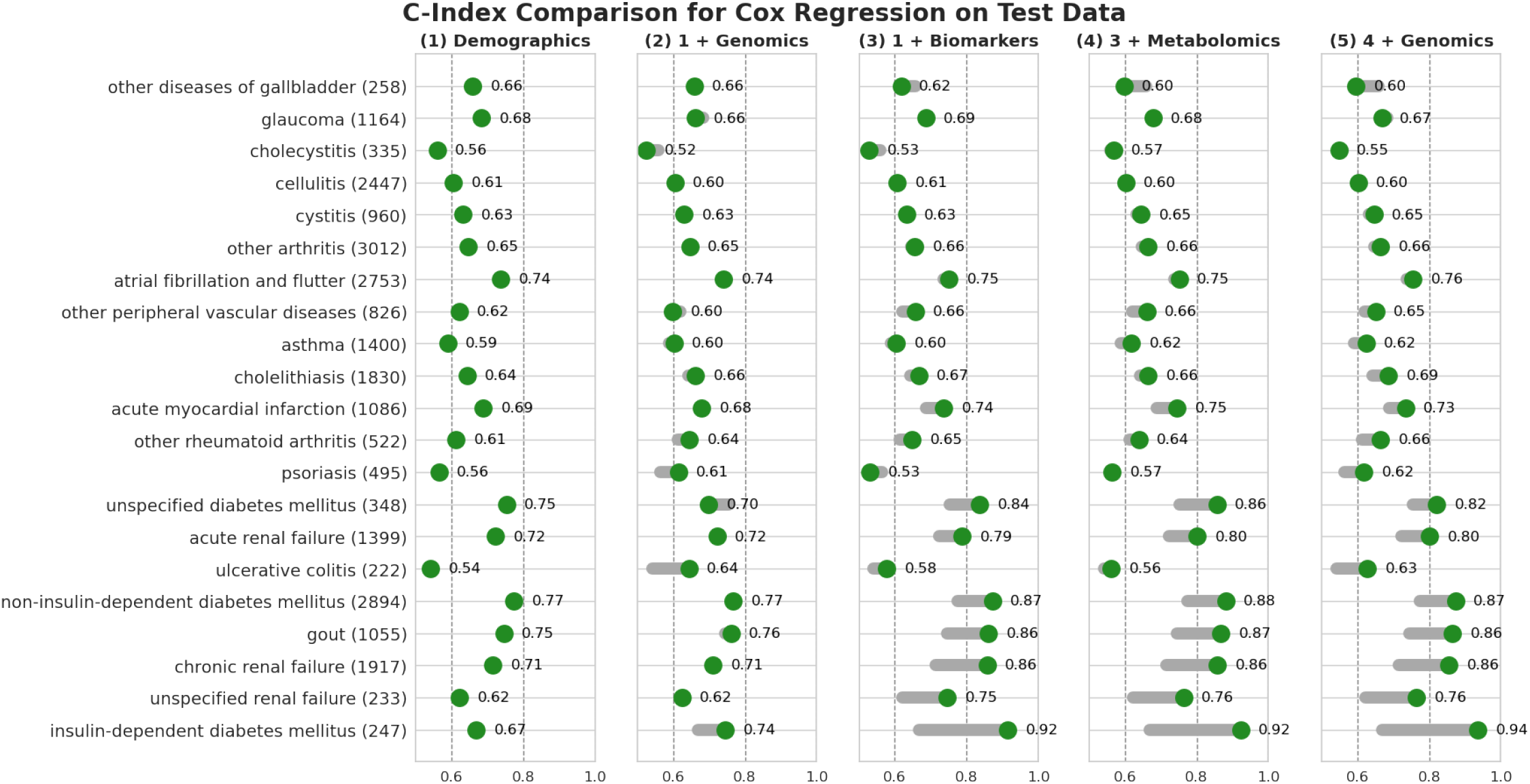
C-index comparison of Cox regression models trained on different subsets of the data. Green dots represent the C-index on the held-out test set for each disease, achieved by a model trained on the indicated group of features. Solid bars represent the change in performance from the baseline demographics model (first column). The numbers in the parentheses by the disease names are the number of incident cases for each disease in the dataset.

Adding the genomics feature to the baseline model results in improved C-indices for some of the disease models. The C-index of the psoriasis model increases from 0.56 to 0.61, the ulcerative colitis model from 0.54 to 0.64, and the insulin-dependent diabetes mellitus model from 0.67 to 0.74. For most diseases, however, the C-indices remain unchanged or have very small C-index increments when compared to the demographics-only model. We observe a decrease in C-indices for the models of cholecystitis and unspecified diabetes mellitus.

By training models with demographic and biomarker data, we observe a larger gain in C-indices for more disease models. For diabetes (insulin-dependent and non-insulin-dependent), renal failure, and gout, incorporating biomarker information improves the performance of the models. C-indices increase from 0.77 to 0.87 for non-insulin-dependent diabetes, from 0.75 to 0.86 for gout, and from 0.71 to 0.86 for chronic renal failure. For other diseases, the gain in C-index with the incorporation of biomarker data is minimal. In general, these results agree with the finding of the binary classification model. For other diseases of gallbladder, cholecystitis, and psoriasis, incorporating biomarkers decreases the C-indices of the resulting models.

Additionally, incorporating genomics and metabolomics data on top of demographics and biomarker features only provides marginal C-index improvement for models of a few diseases, such as psoriasis (from 0.64 to 0.66) and cholecystitis (from 0.53 to 0.55) (Figure 3 Column 5). C-indices of most models remain the same, while the model performance of other diseases of gallbladder and unspecified diabetes mellitus decreases. This is also largely in agreement with the results of the binary classification model.

We use the combined dataset of all demographic, genomics, biomarker, and metabolomics features (Figure 3 Column 5) to find the optimal regularization strength of the L1 penalty, evaluated by their C-indices over the validation dataset [8]. We use this process to inspect feature relevance and importance to each disease. Figure 4 shows a sample feature importance plot generated by the hyperparameter search. Omitted features are those with a coefficient value of 0 based on the optimal L1 penalty. For the “other peripheral vascular disease” model, a subset of 40 features have non-zero coefficients after feature selection. For most diseases, among the 288 potential features, there are about 10 to 50 features in the L1 regularized models, with age, sex, and the corresponding PRS of the diseases being the most frequently kept features. The sparsity of significant features based on L1 regularization varies for different diseases, with the final feature count ranging from 2 (cystitis) to 95 (other arthritis).

**Fig. 4:**
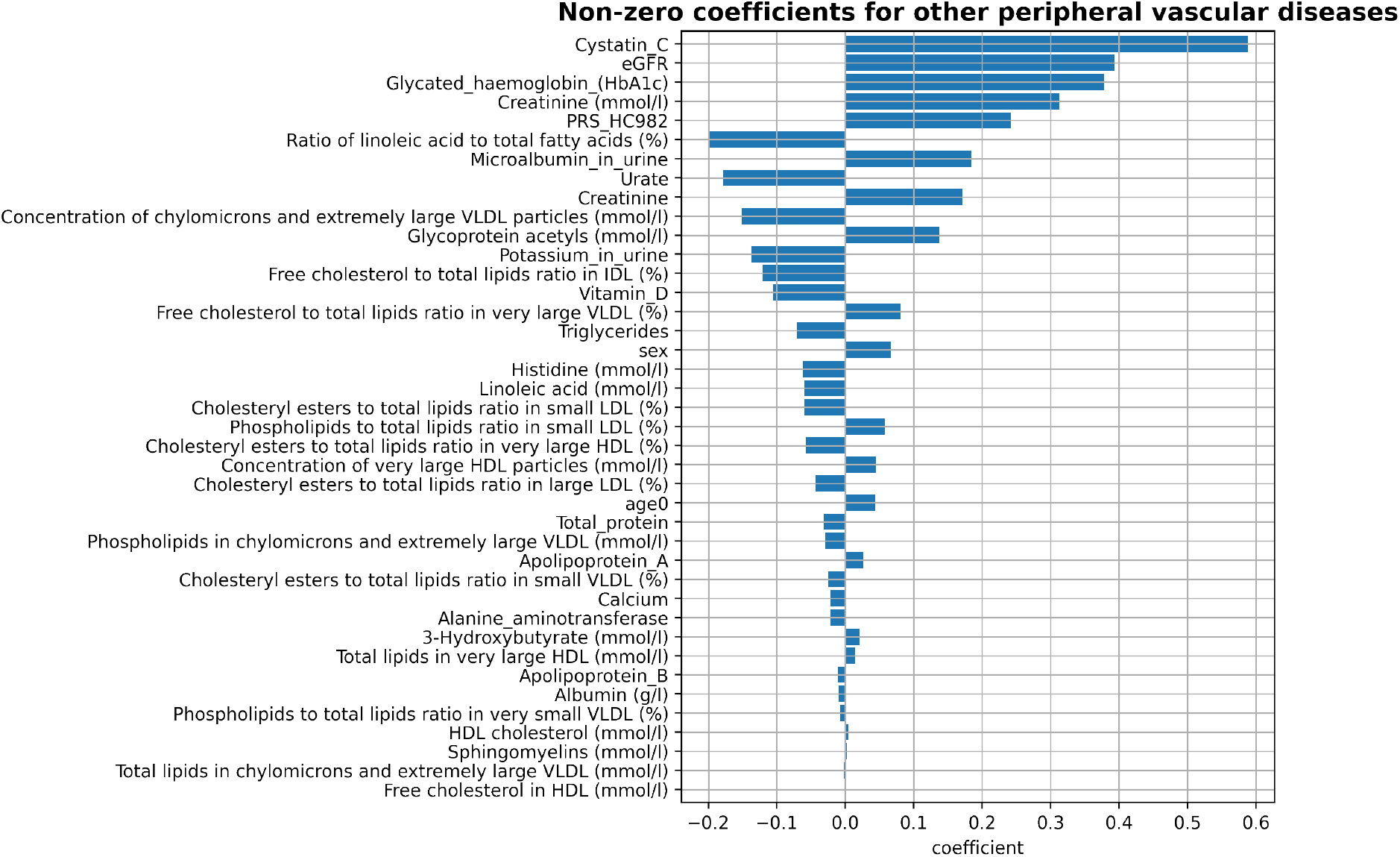
Sampled feature importance plot for other peripheral vascular diseases. Any omitted feature indicates a coefficient value of 0 from L1 regularization.

Finally, we train all L1 regularization Cox models with the full training dataset and evaluate the model performance on the previously unseen test dataset. For some diseases, such as diabetes and renal failure, the corresponding models have a test C-index above 0.8. Most of the disease-specific models have C-indices above 0.6. See Figure B3 for the final training and test C-index of each disease-specified model.

Using the model we trained, we can predict the risk trajectories of each disease for each individual in the dataset (Figure 5). Observations in the figure are randomly sampled from the test dataset of chronic renal failure. 2 positive and 2 negative observations are sampled. Here, a positive sample means a participant of the study who acquired the disease during the period of the study. A negative sample indicates a participant who did not acquire the disease or did not acquire it during the period in which the participant continued to be part of the study (right-censored). Each line indicates a predicted trajectory of disease onset probability for an individual, with the legend marking their actual test results for the disease.

**Fig. 5:**
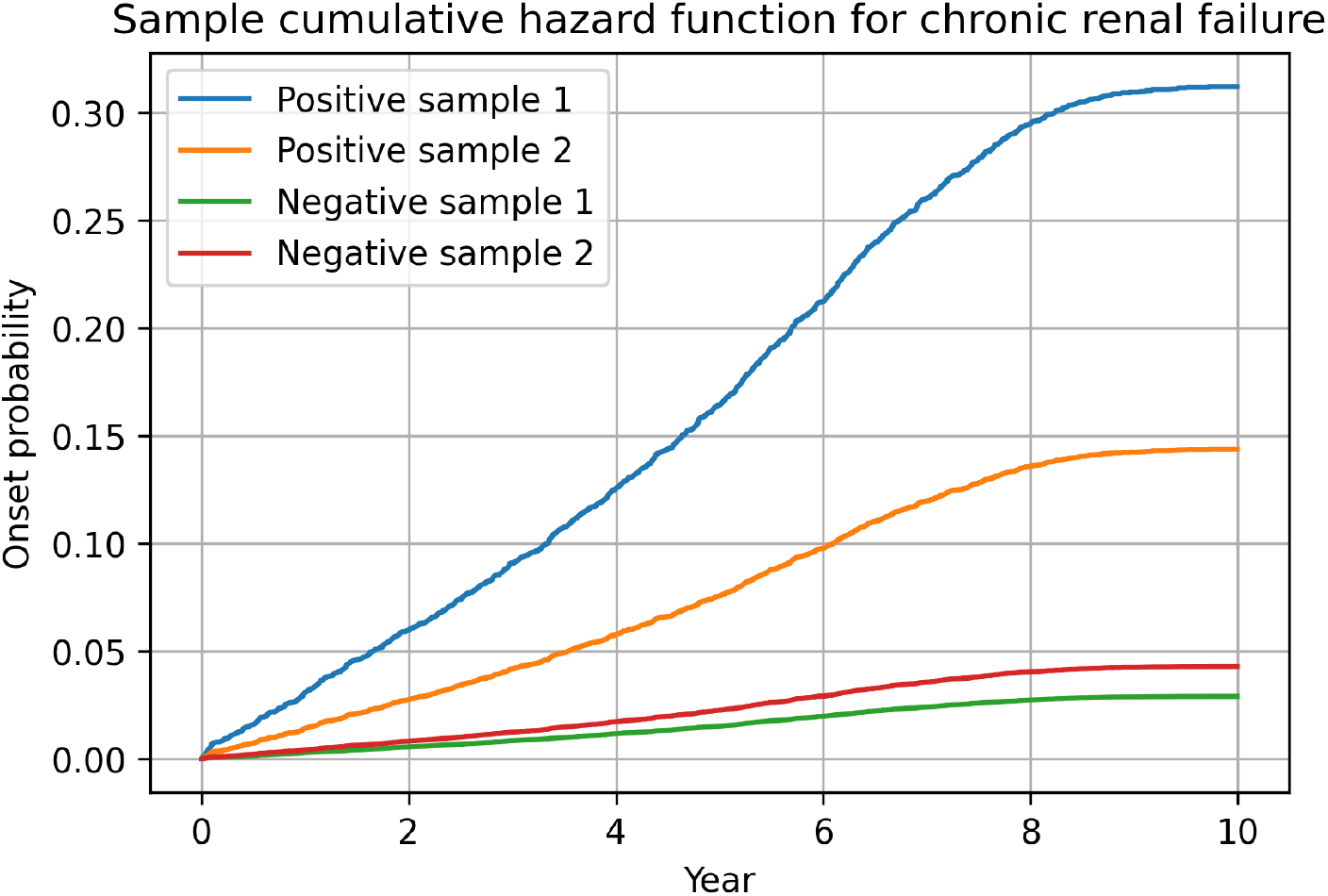
Predicted risk trajectories of chronic renal failure of 4 participants in the test dataset. Each line represents a select individual from the test dataset and their estimated hazard trajectory throughout the years. Here, a positive sample means a participant of the study who acquired the disease during the period of the study. A negative sample indicates a participant who did not acquire the disease or did not acquire it during the period in which the participant continued to be part of the study (right-censored).

## 3 Discussion

Our analysis of the prediction models highlights the strengths and weaknesses of different classifiers. XG Boost took the least amount of time to fit each model — an average of 20 seconds using default parameters versus 156 for ADA Boost. Meanwhile, ADA Boost yielded the sparsest and most interpretable models, selecting on average 20% the number of non-zero features selected by Lasso (Table A1). Lasso regression took the most time to train and tune — a total of 208 seconds per disease — but gave the best results once the proper imputation, scaling, and regularization paths were followed. Notably, the Lasso models had the highest AUC on the test set for 18 out of 22 diseases, with median AUC 0.012 points (1.7%) higher than the ensemble classifiers (Table 1). Interestingly, the Multi-Layer Perceptron showed performance in line with Lasso for certain diseases, including an improvement in Test AUC of 1% for myocardial infarction. Overall, this work builds upon many previous findings of linear models providing a dominant performance on the task of disease prediction.

In addition to the binary classification models, we conduct survival analysis using the Cox proportional hazard model. We compare the contribution of different feature types toward model performance using C-indices and search for relevant features for each disease with L1 regularization. The survival analysis provides additional information about the disease onset time and the trajectories of risk scores as patients age. The feature type analysis and model training results of the survival analysis model largely confirm the findings of the binary classification models.

By evaluating the contributions of demographic, genomics, metabolomic, and biomarker data over the classification and survival analysis model, we can identify diseases with strong genetic or metabolomic components and observe the incremental improvements achieved by adding various feature types. We assessed marginal improvements in model performance from combining genomic and metabolomic data into a multi-omics model, particularly for diseases with both a genetic and a metabolomic component. Finally, for most diseases with a metabolomic component, we observed no significant improvement in model performance upon expanding the set of 35 biomarkers with all 249 metabolites. This corroborates the findings of Sinnott-Armstrong et al. in [1] regarding the utility of these 35 particular blood and urine biomarkers in the UK Biobank. In addition, this invites the question of whether a metabolomics panel could be a substitute for the standard lab panels used for disease risk assessment. Indeed, the comparison of adding metabolomic vs biomarker data on top of demographic and genomic data for disease prediction in Figure A1 shows equivalent model performance for every disease except type 2 diabetes, which performs worse by 0.03 AUC. The time-to-event models in Figure B2, however, display a larger loss of performance from replacing clinical biomarkers with metabolite levels.

A limitation of our work is the low disease onset count in our dataset. For example, the results we generated for diseases with particularly low case counts, such as those from the binary classification models of ulcerative colitis (*n* = 916) and the survival model of unspecified renal failure (*n* = 233), will be more vulnerable to instability. As the cohort of the Biobank dataset ages, more patients may contract our diseases of interest, but this also means that we may have a clearer picture of their health status later in life. Therefore, continual investigations of the health status of patients in the dataset may help alleviate the imbalance dataset problem.

Our models are primarily based on data from the white British population within the UK Biobank dataset. Consequently, our findings may not apply to other nationalities or ethnic groups. Prior studies have shown that similar models developed within one ethnic group have limited transferability for other groups [9]. Therefore, future work with our models could explore the effect of our model and training methods on other populations. In addition, providing measures of uncertainty for the Multiomic Risk Scores from our classification models could help guide the implementation of these models in. Finally, we recommend further investigation into the diseases for which we identified small multi-omic benefits and the development of better methods for fusing the information from the different “omic” datasets. One such method introduced by Ding, Li, Narashimiran, and Tibshrani in [10] is called “cooperative learning”. In our multiview setting, this approach would encourage the predictions from our demographic, metabolomic, and genomic models to agree. This would also extend to the further incorporation of proteomic and other multi-modal datasets into our experiments.

## 4 Methods

### Data

We use four distinct datasets obtained from the UK BioBank, covering demographics, genomics, metabolomics, and clinical biomarkers. The full dataset includes information from 488,377 individuals of various origins within the UK BioBank. Demographic features include age, sex, and Body Mass Index (BMI). The genomics data for these individuals is precomputed into a Polygenic Risk Score (PRS) for each disease, calculated without the incorporation of demographic features in a prior study using the BASIL algorithm [9]. The clinical biomarkers dataset includes a curated set of 35 blood and urine biomarkers, selected by previous studies based on their relevance [1]. The metabolomics dataset comprises a larger set of 249 biomarkers in the blood, available for a subset of 87,465 White-British individuals.

Our disease prediction models use this set of 87,465 White-British individuals for training and testing. Additionally, a set of 29,089 non-White British individuals is used to train/test different strategies for incorporating the PRS without touching the heldout test set. See [11] for the classification of ancestry in the UK Biobank. The set of 22 diseases analyzed with our binary classification models includes diabetes, asthma, renal failure, and a range of cancers. For survival analysis using the Cox proportional hazard model, the same datasets serve as potential features. Additionally, an age-of-onset dataset specific to each disease facilitates the prediction task. The diseases under consideration span a spectrum, including asthma, type-1 and type-2 diabetes, arthritis, cellulitis, cholelithiasis, atrial fibrillation and flutter, myocardial infarction, renal failure, cystitis, gout, glaucoma, psoriasis, peripheral vascular diseases, rheumatoid arthritis, ulcerative colitis, cholecystitis, and diseases affecting the gallbladder.

### Binary Disease Onset Classification

We applied several tree-based ensemble models from *sklearn*.*ensemble* and *xgboost*.*XGBClassifier*, logistic regression models from *sklearn*.*linear model* with various regularizations, and some deep learning classifiers from *sklearn*.*neural network*. Each model was tuned using a 1-fold cross-validation with a predetermined train-validation split. To ensure consistency with the development of the PRS scores for each disease/biomarker [9], the same 70/10/20 train/validation/test split of white British individual data was used.

For the ensemble models, we allowed the number of estimators to vary between 5 and 200. For the linear models, we allowed the regularization strength to vary logarithmically between 1e-4 and 1e4. For the multi-layer perceptron, we explored using 2 hidden layers with between 5 and 50 nodes each. Most ensemble models selected 10 estimators, while the selected hyperparameters for the MLP and Linear classifiers varied more. In particular, it was important to correctly tune the linear models to achieve the best overall performance. This process was made more efficient by following a regularization path which warm-started each fit with the previous one, implemented natively in *sklearn*.*linear model*.*LogisticRegressionCV*.

Next, feature importance analysis was performed for the ensemble and linear classifiers to judge model quality and perform feature selection. Relative feature importance for each model was extracted via YellowBrick’s *model selection*.*FeatureImportances* module. For our logistic regression models, we added standard scaling as a preprocessing step in order to properly compare coefficients.

### Survival Analysis

The same predefined train-validation-test split (70/10/20) was used to train and fine-tune the model. Data is pre-processed by feature-wise normalization and median imputation (replacing missing data with the median). Samples with a disease onset time earlier than the data collection time are left-censored. Note that we left-censor the data by disease; data samples that are excluded for one type of disease may be used to train models of other diseases. This means that the set of datasets used to train and evaluate each model is unique.

By left-censoring, we ensure that only incident cases of each disease are left in the dataset. Figure 3 shows the number of incident cases for each disease. The number of cases ranges from 222 (ulcerative colitis) to 3012 (other arthritis). For most diseases, the number lies between 500 to 2000.

We first train models for each disease with different combinations of data types on the training dataset. The data type groups we explore include (1) demographics; (2) demographics and genomics; (3) demographics and biomarkers; (4) demographics, biomarkers, and metabolomics; and (5) all 4 feature types (see Figure 3). We compare the C-index of the five models for each disease. We then used the validation dataset to conduct a hyperparameter search over *α*, the L1 regularization strength [8]. For each disease, we generate and evaluate the feature importance plots (Figure 4) with the *α* that achieves the highest C-index. We use the training dataset for our final model training. The models are then evaluated on the test dataset.

We use *CoxnetSurvivialAnalysis* in the Python *scikit* − *learn* library, which is a Python re-implementation of the *Coxnet* implementation in *R*, throughout our model development and analysis process [12].

## Supplementary information

The code for our model development and data analysis is hosted on a GitHub repository at https://github.com/rivas-lab/multiomics-ukb.

## Acknowledgements

M.A.R. is in part supported by National Human Genome Research Institute (NHGRI) under award R01HG010140, and by the National Institutes of Mental Health (NIMH) under award R01MH124244 both of the National Institutes of Health (NIH).

Some of the computing for this project was performed on the Sherlock cluster. We would like to thank Stanford University and the Stanford Research Computing Center for providing computational resources and support that contributed to these research results. The content is solely the responsibility of the authors and does not necessarily represent the official views of the funding agencies; funders had no role in study design, data collection and analysis, decision to publish, or preparation of the manuscript.

This research has been conducted using the UK Biobank Resource under Application Number 24983, “Generating effective therapeutic hypotheses from genomics and hospital linkage data” (http://www.ukbiobank.ac.uk/wp-content/uploads/2017/06/24983-Dr-Manuel-Rivas.pdf). Based on the information provided in Protocol 44532, the Stanford IRB has determined that the research does not involve human subjects as defined in 45 CFR 46.102(f) or 21 CFR 50.3 (g). All participants of the UK Biobank provided written informed consent (information available at https://www.ukbiobank.ac.uk/2018/02/gdpr/).

## Appendix A Binary Classification Models

**Table A1:** Comparison across different models of the number of features with non-zero importance for every disease.

| Model | ADA Boost | Lasso | XG Boost |
| --- | --- | --- | --- |
| arthritis (nos) | 67 | 164 | 121 |
| asthma | 26 | 34 | 178 |
| atrial flutter | 32 | 72 | 165 |
| bowel cancer | 5 | 10 | 112 |
| breast cancer | 7 | 11 | 144 |
| cellulitis | 7 | 22 | 131 |
| cholelithiasis | 67 | 154 | 173 |
| chronic renal failure | 54 | 157 | 167 |
| cystitis | 5 | 5 | 114 |
| diabetes | 26 | 190 | 175 |
| female genital tract cancer | 7 | 44 | 79 |
| gall stones | 32 | 74 | 174 |
| glaucoma | 3 | 59 | 95 |
| gout | 32 | 65 | 162 |
| irritable bowel syndrome | 5 | 61 | 183 |
| lung cancer | 33 | 81 | 186 |
| male genital tract cancer | 6 | 268 | 111 |
| myocardial infarction | 35 | 288 | 181 |
| non-melanoma skin cancer | 3 | 14 | 125 |
| osteoarthritis | 29 | 288 | 167 |
| peripheral vascular disease | 42 | 288 | 173 |
| prostate cancer | 4 | 12 | 136 |
| psoriasis | 2 | 57 | 128 |
| renal failure | 32 | 162 | 171 |
| skin cancer | 4 | 14 | 184 |
| ulcerative colitis | 3 | 49 | 83 |
| Avg Number of Features | 22 | 103 | 147 |

**Table A2:**
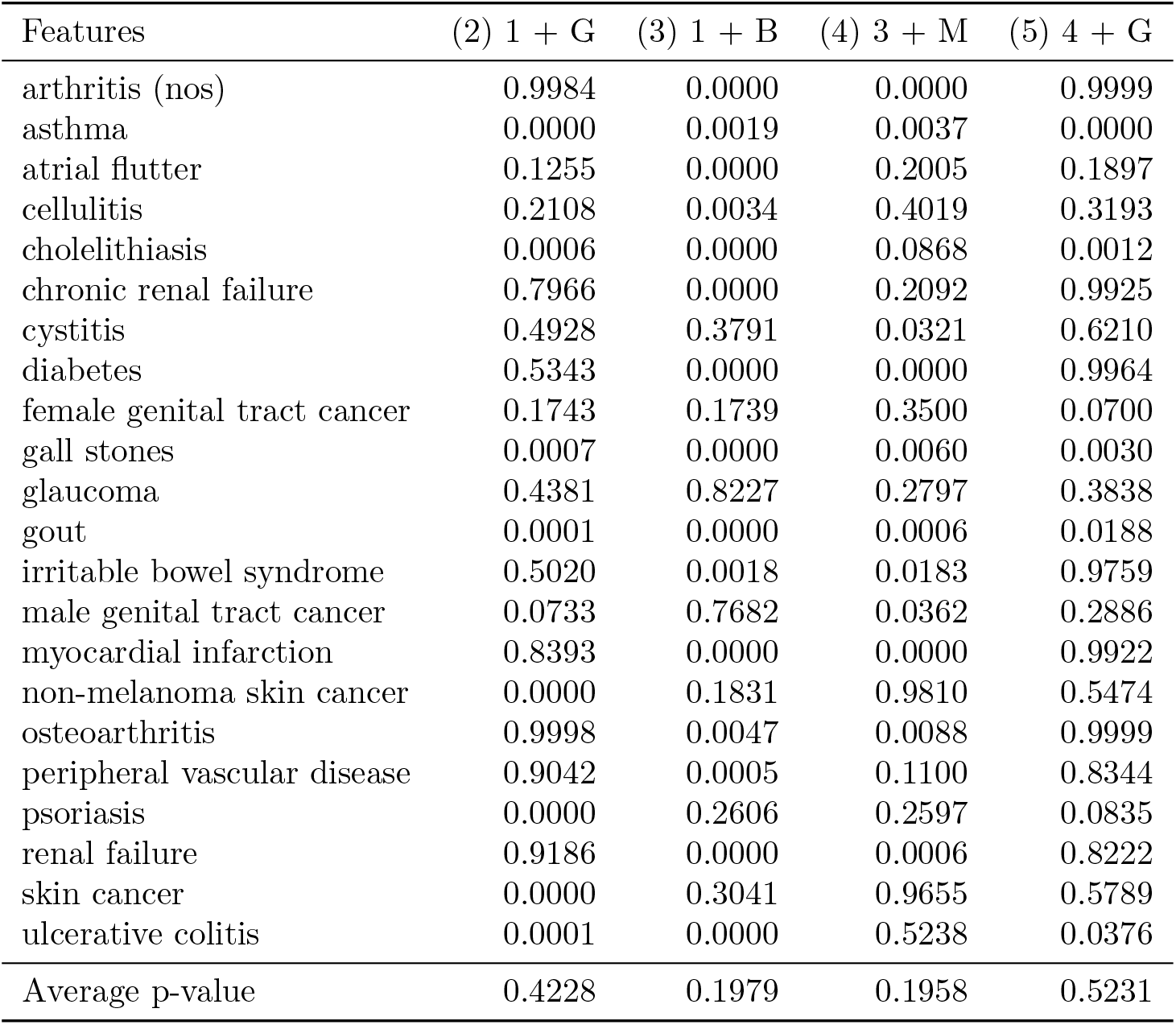
P-value comparison of the ROC curves from consecutive lasso regression models trained on different subsets of the data. For each column in Figure 2, the model predictions on the test set were compared to the predictions from the prior model, referenced by column number. To assess the significance of the observed delta AUC, a permutation test is performed on the paired ROC curves. The p-value is the empirical probability of observing a larger improvement in AUC on a shuffled data set.

| Features | (2) 1 + G | (3) 1 + B | (4) 3 + M | (5) 4 + G |
| --- | --- | --- | --- | --- |
| arthritis (nos) | 0.9984 | 0.0000 | 0.0000 | 0.9999 |
| asthma | 0.0000 | 0.0019 | 0.0037 | 0.0000 |
| atrial flutter | 0.1255 | 0.0000 | 0.2005 | 0.1897 |
| cellulitis | 0.2108 | 0.0034 | 0.4019 | 0.3193 |
| cholelithiasis | 0.0006 | 0.0000 | 0.0868 | 0.0012 |
| chronic renal failure | 0.7966 | 0.0000 | 0.2092 | 0.9925 |
| cystitis | 0.4928 | 0.3791 | 0.0321 | 0.6210 |
| diabetes | 0.5343 | 0.0000 | 0.0000 | 0.9964 |
| female genital tract cancer | 0.1743 | 0.1739 | 0.3500 | 0.0700 |
| gall stones | 0.0007 | 0.0000 | 0.0060 | 0.0030 |
| glaucoma | 0.4381 | 0.8227 | 0.2797 | 0.3838 |
| gout | 0.0001 | 0.0000 | 0.0006 | 0.0188 |
| irritable bowel syndrome | 0.5020 | 0.0018 | 0.0183 | 0.9759 |
| male genital tract cancer | 0.0733 | 0.7682 | 0.0362 | 0.2886 |
| myocardial infarction | 0.8393 | 0.0000 | 0.0000 | 0.9922 |
| non-melanoma skin cancer | 0.0000 | 0.1831 | 0.9810 | 0.5474 |
| osteoarthritis | 0.9998 | 0.0047 | 0.0088 | 0.9999 |
| peripheral vascular disease | 0.9042 | 0.0005 | 0.1100 | 0.8344 |
| psoriasis | 0.0000 | 0.2606 | 0.2597 | 0.0835 |
| renal failure | 0.9186 | 0.0000 | 0.0006 | 0.8222 |
| skin cancer | 0.0000 | 0.3041 | 0.9655 | 0.5789 |
| ulcerative colitis | 0.0001 | 0.0000 | 0.5238 | 0.0376 |
| Average p-value | 0.4228 | 0.1979 | 0.1958 | 0.5231 |

**Fig. A1:**
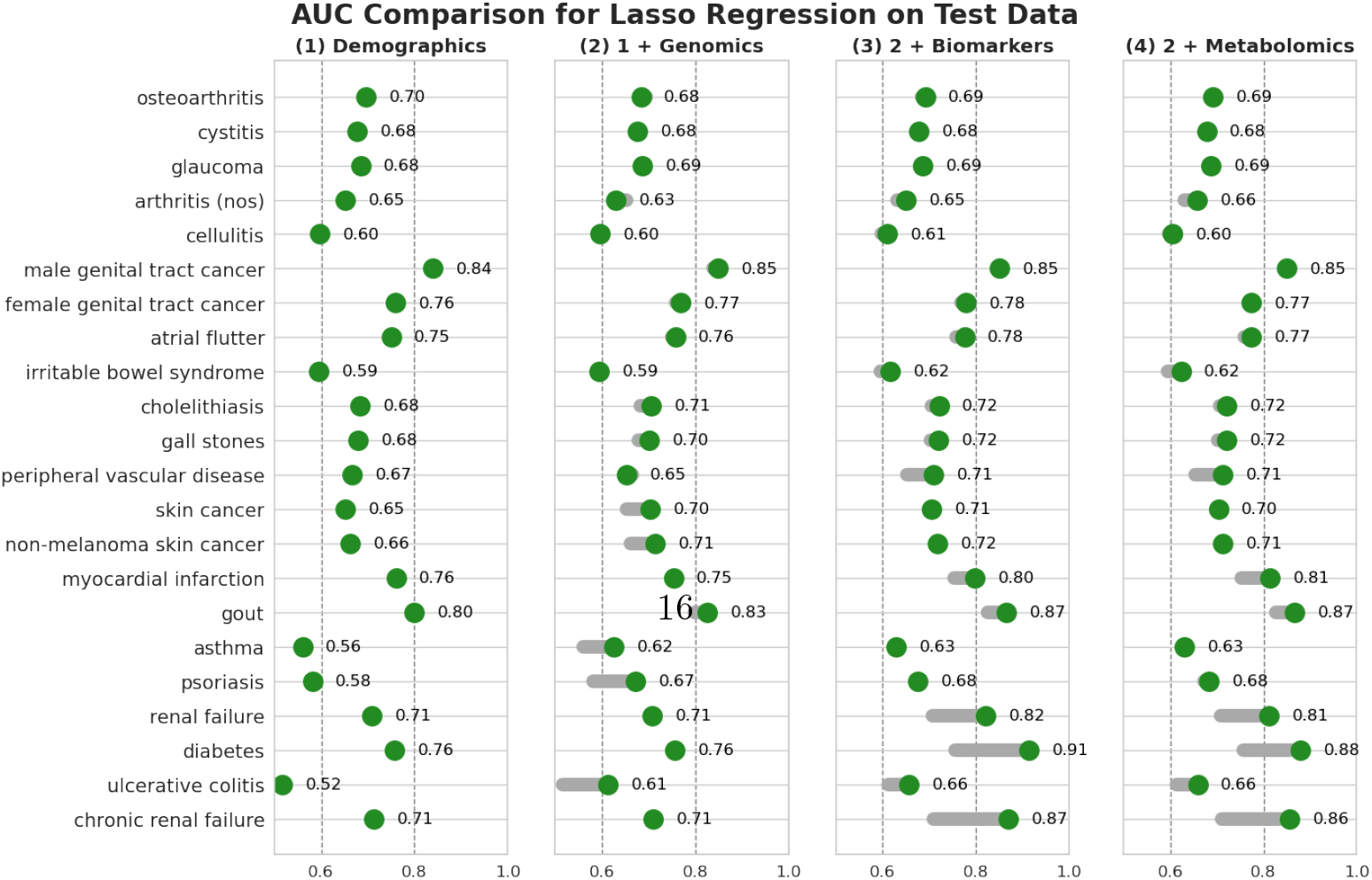
Comparison of the test AUC of lasso regression models trained on different subsets of the data. Green dots represent the AUC (area under the ROC curve) on the test set for each disease, achieved by a model trained on the indicated group of features. Solid bars represent the change in performance from the baseline demographics model (first column).

## Appendix B Survival Models

### B.1 Model Evaluation

**Fig. B2:**
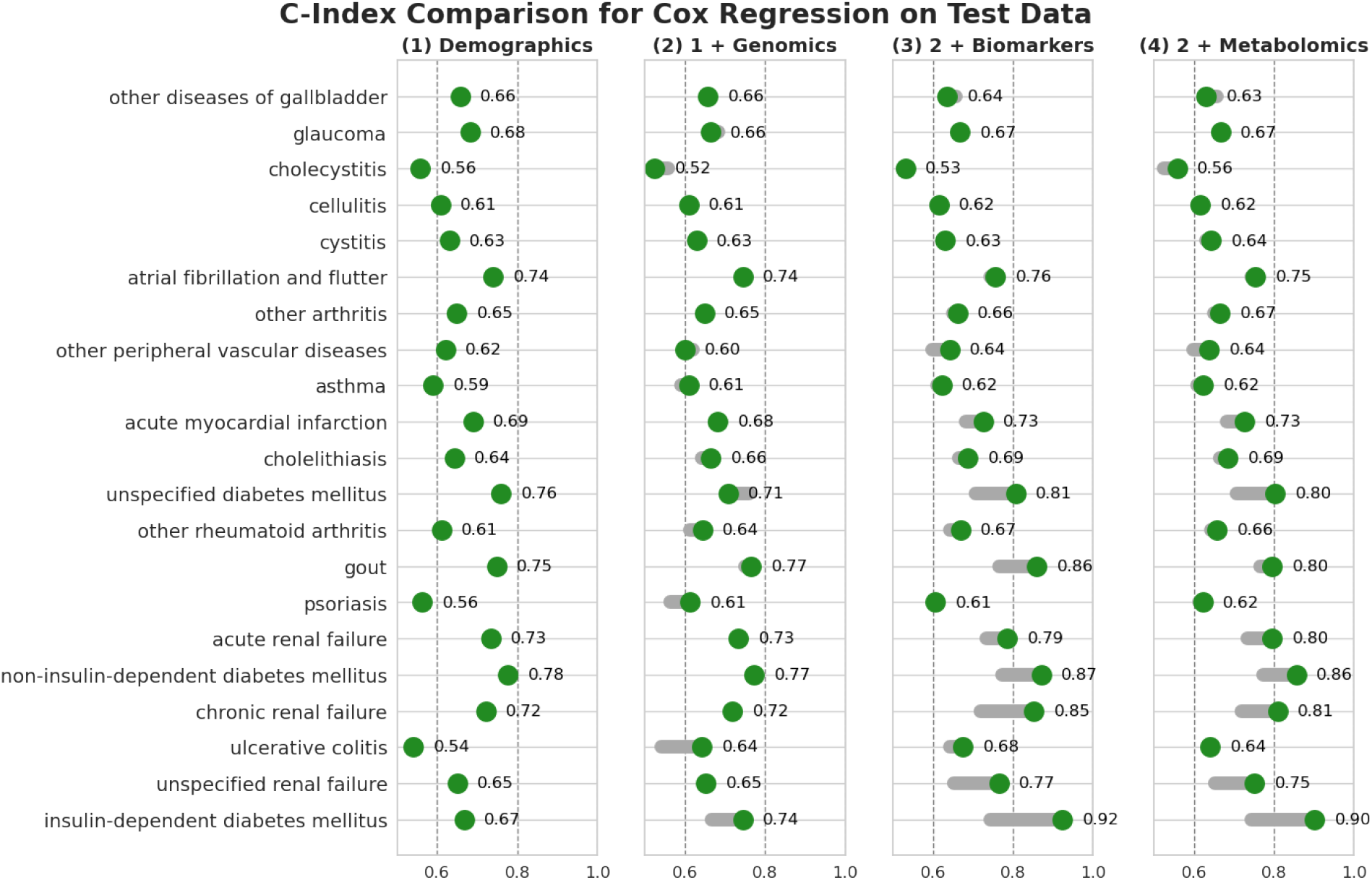
Comparison of the C-index of Cox regression models trained on different subsets of the data. Green dots represent the C-index on the test set for each disease, achieved by a model trained on the indicated group of features. Solid bars represent the change in performance from the baseline demographics model (first column).

**Fig. B3:**
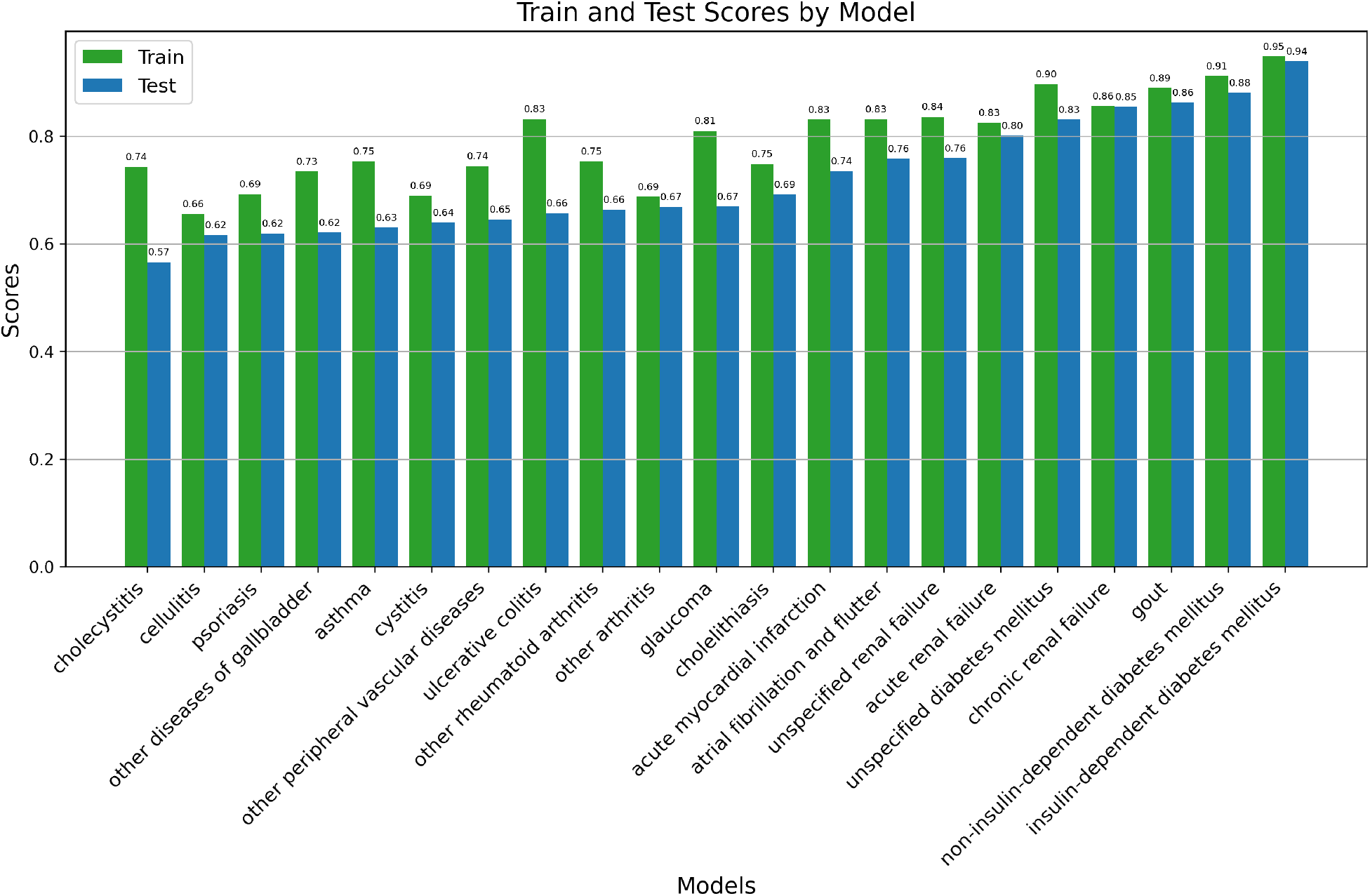
Training and test C-index comparison for all diseases.

